# Tac1 Deficiency Reduces the Severity of Enteric Bacterial Infection

**DOI:** 10.1101/2025.05.27.656414

**Authors:** Elliot Lloyd, Micheal Cremin, Kristina Sanchez, Jungjae Park, Melanie Gareau, Colin Reardon

## Abstract

**Background:** Infection with enteric bacterial pathogens continues to cause significant morbidity and mortality throughout the world. These pathogens include enterohemorrhagic and enteropathogenic *Escherichia coli*, which transit the intestinal tract, efface microvilli, and attach firmly to intestinal epithelial cells predominantly in the colon. Investigation of these human-adapted pathogens has been greatly aided by mouse models of infection. The mouse-adapted attaching and effacing pathogen *Citrobacter rodentium* utilizes many similar mechanisms of pathogenesis, including the use of a type III secretion system, and virulence factors encoded in a locus of enterocyte effacement. Although this model has allowed for assessing the complexity of the host response, the complex interplay between the nervous and immune systems in response to infection remains incomplete.

**Methods:** We assessed the role of sensory neurotransmitters encoded by the *Tac1* gene in the host response to *C. rodentium*.

**Results:** *Tac1*-deficient mice had significantly reduced pathogen shedding and colonic bacterial burden, accompanied by decreased expression of inflammatory cytokines and chemokines. In accordance with reduced chemokine production, we observed reduced colonic recruitment of specific immune cell populations in *Tac1-/-* compared to WT mice.

**Conclusions:** Sensory neuropeptides regulate key aspects of enteric bacterial infection and may serve as unique targets in the treatment of enteric disease.

## Introduction

The ability to mount immune responses to pathogens while limiting the potential for aberrant or excessive inflammation that can harm the host is critical to protective responses. In the intestinal tract, the mucosal immune system distinguishes between noxious substances, potential pathogens, and substances derived from the diet and commensal microbiota. Therefore, processes that regulate immune responses in mucosal tissues, such as the colon, are critical to health. Our understanding of these processes has been greatly aided by the mouse-adapted pathogen, *Citrobacter rodentium* [1]. Protective host responses include the activation of innate lymphoid cells (ILC) and tissue-resident macrophages, the recruitment of neutrophils, monocytes, and eventually T-cells. These responses are coordinated throughout infection to culminate in the eventual eradication through B-cell-produced immunoglobulins. Although cytokines and chemokines produced by immune cells are required to enable this coordination, the nervous system’s production of neurotransmitters can alter the trajectory of an immune response and thus the course of disease.

The intestine is replete with intrinsic innervation, which comprises the enteric nervous system (ENS), and extrinsic innervation, which originates in the dorsal root and sympathetic chain ganglia [2]. While the ENS controls many physiological functions of the intestine, extrinsic innervation can also influence these processes. Detection of noxious stimuli such as excessive heat, inflammatory products, or bacterial components can be achieved by specialized sensory neurons expressing the polymodal nociceptor TRPV1 [3, 4]. Activation of TRPV1, and consequently these neurons, can cause the localized release of peptide neurotransmitters such as Substance P (SP) [5–7]. During enteric bacterial infection, we and others have demonstrated that TRPV1+ innervation coordinates the host immune response to bacterial enteric infection [8, 9]. Using pharmacological and genetic approaches, we have also shown that Substance P (SP) receptor signaling fine-tunes the host response, with unique cell types expressing this receptor controlling T-cell recruitment and IFNγ production [10]. These findings are in keeping with the defined roles of this transmitter during neurogenic inflammation, where SP increases blood vessel dilation and permeability, and aids the recruitment of immune cells by increasing the expression of adhesion molecules by endothelial cells [11–13]. It is also now appreciated that SP can exert biological effects, such as mast cell degranulation, through activation not only of SP receptors but through Mas-related G protein receptors [14]. As a result, the biological effect of SP in the intestine is of considerably greater complexity than previously appreciated. This complexity extends to the source of substance P in the intestine, with expression in extrinsic sensory neurons, putative interneurons, motor neurons of the ENS [15, 16] and enteroendocrine cells [17–19]. The unique positioning of these cells in the intestine, coupled with the potential for SP-mediated activation of responding cells through at least one other receptor, suggests potentially diverse roles of SP in regulating host responses.

With this in mind, we sought to characterize the response to enteric bacterial infection with *C. rodentium* using mice with global disruption of the *Tac1* gene that encodes SP and neurokinin A among others. Using this approach, we found significantly reduced bacterial shedding in the feces and reduced colonic bacterial burden in *Tac1^-/-^* mice compared to wild-type (WT). Consistent with these reductions in *Tac1^-/-^* mice, infection-induced colonic histopathology was significantly attenuated relative to infected WT mice. Given the importance of SP to intestinal physiology, Ussing chambers were used to determine the effect of *Tac1* deficiency on baseline and induced ion secretion, conductance and permeability. In assessing the innate and adaptive immune responses during infection, significant reductions in TNFα and select chemokines that aid recruitment of neutrophils and T-cells were observed in Tac1-/- mice compared to WT. Significantly reduced IFNγ, and expression of IFNγ-dependent genes such as inducible Nitric oxide synthase (*Nos2*), were also found in infected *Tac1^-/-^* compared to WT mice. These changes in chemokines and commonly elicited T-cell cytokines in response to *C. rodentium* were matched by reduced recruitment of these cells into the infected colons of *Tac1^-/-^* mice. With the noted reduction in specific T-cell responses*, in vitro* Th1 differentiation was performed, and no intrinsic defects in generating these responses were revealed. Together, these data demonstrate that the Tac1 gene coordinates specific aspects of the host response to enteric bacterial pathogens, including *C. rodentium*.

## Methods

### Animals

C57BL/6J, *Tac1-/-*, mice were purchased from Jackson Laboratories and used to establish a breeding colony at UC Davis. Male and female mice aged 7-12 weeks were used for experiments. All animals had *ad libitum* access to food and water, and all procedures were approved by the UC Davis Institutional Animal Care and Use Committee (IACUC).

### *Citrobacter rodentium* infection and enumeration of bacteria in feces and colon

*C. rodentium*, strain DBS100, (generously provided by Dr. Andres Baumler) was grown on MacConkey agar plates (37°C overnight), followed by overnight culture in Luria broth (LB) at 37°C without shaking. Mice were gavaged with LB or *C. rodentium* (10^8^ CFU (colony-forming units)). Enumeration of bacteria was performed with fecal pellets of an approximately 1 cm long piece of distal colon. Samples were placed into a pre-weighed microcentrifuge tube, and weight determined, followed by homogenization in a TissueLyser (Qiagen). Homogenates were serially diluted and plated on MacConkey agar, incubated (16h, 37°C), and counted, to allow for CFU per gram of sample to be calculated.

### Distal colonic motility – fecal output

Distal colonic motility was measured in uninfected WT and *Tac1-/-* mice by assaying the number of fecal pellets, in a 20-minute period. Pellet weight was also determined without dehydration.

### Quantitative real-time PCR

Colon tissue samples were collected in TRIzol (Invitrogen) and frozen (-80°C). Tissues were homogenized using stainless steel beads in a TissueLyser (Qiagen), and RNA extracted. Synthesis of cDNA using the isolated RNA was performed with the iSCRIPT reverse transcriptase kit (Bio-Rad, Hercules, CA, USA), followed by qPCR analysis using a QuantStudio6 (Thermo Fisher Scientific, Waltham, MA, USA) machine with the primer pairs indicated **(Table 1)** [20].

**Table 1.**
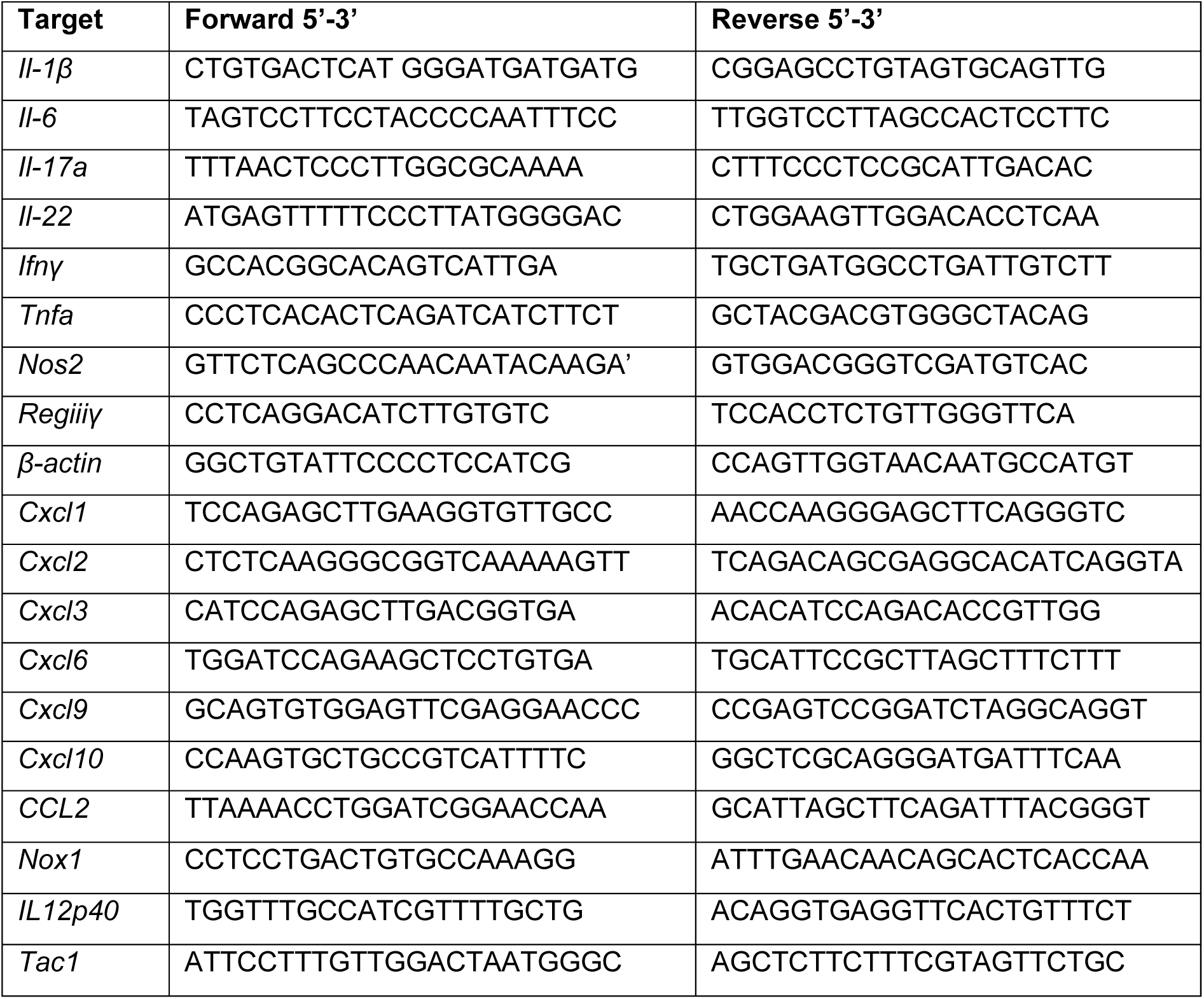
Primer sequences.

### *In vitro* T-cell differentiation and ELISA

T-cells were enriched by negative selection using the iMAG system (BD Biosciences, Milpitas, CA, USA). In brief, single-cell suspensions from the mesenteric lymph nodes were incubated with Fc block (anti-CD16/32) for 20 minutes on ice, followed by incubation in a cocktail of biotinylated antibodies directed against Ter119, CD8, CD11b, CD11c, B220, CD317, and NK1.1. After a 30-minute incubation on ice, labelled cells were washed, and mixed with SA-iMAG magnetic beads (Cat# 557812, BD Biosciences). These samples were placed within the magnetic field of the iMAG magnet, allowing for recovery of the non-labelled cells in the supernatant. Recovered negatively enriched CD4+ T-cells were plated on round bottom 96-well plates at a density of 25,000 cells/well. Wells were pre-treated with 2 µg/mL anti-CD3 (Cat# 40-0032 Tonbo Biosciences San Diego, CA) in PBS for 2 h at 37°C and 5% CO_2_. Cells were cultured with or without 0.5 µg/mL anti-CD28 (Cat# 40-0281 Tonbo Biosciences) and in Th1 skewing culture conditions. This consisted of 1 µg/mL anti-murine IL-4 (Cat# 16-7041 Invitrogen Carlsbad, CA), 5 ng/mL murine IL-2 (Cat# 212-12 Peprotech Cranbury, NJ), and 10 ng/mL murine IL-12p40 (Cat# 210-12 Peprotech). Cells were incubated 37°C 5% CO_2_ for 96 h and stimulated with 1X Cell Stimulation Cocktail (phorbol 12-myristate 13-acetate) (eBioscience, San Diego CA) for 4 h. Supernatant was frozen at -20°C until assayed by ELISA. Untreated 96-well flat-bottom plates were coated overnight at 4°C with capture antibodies for IFNγ (Cat# 88-7314-22, Invitrogen), following the manufacturer’s protocols. Plates were read on a Synergy HTX Multi-Mode Plate Reader (BioTek, Winooski, Vermont, USA) using the Gen5 application at 450 nm and 570 nm, with a standard curve determined using a 4-parameter logistic fit.

### Lamina propria lymphocyte isolation

In brief, colons were removed, opened longitudinally rinsed in HBSS, and cut into 1 cm long segments. Epithelial cells removed by gentle agitation (200 RPM) in HBSS supplemented with 10 mM HEPES, 5 mM EDTA and 5% FBS and discarded. Single-cell suspensions were produced using a lamina propria dissociation kit (Miltenyi Biotec, Gaithersburg, MD) with a gentleMACS tissue dissociator following the manufacturer’s protocol. The resulting single-cell suspension was passed through a 100 µm strainer, washed extensively, and subjected to staining.

### Flow cytometry

Standard flow cytometry staining protocols were followed, where cells were counted by hemocytometer with trypan blue exclusion and incubated with Fc block (anti-CD16/32, 10 µg/ml, Tonbo Biosciences, San Diego, CA) for 25 minutes on ice before incubation with antibody cocktail **(Table 2)** for an additional 30 minutes. Viability was determined using live/dead fixable aqua according to manufacturer’s instructions (ThermoFisher, Waltham MA). Cells were fixed using BD Cytofix (BD Biosciences, Franklin Lakes NJ) for 25 minutes on ice. All flow cytometry data was acquired on a LSRII (BD Biosciences, Franklin Lakes NJ) using DIVA software, with analysis using FlowJo (Becton Dickinson, Eugene OR). Cells were defined based on surface marker expression: T cells (CD45^+^ CD3^+^CD4^+/-^), neutrophils (CD45^+^ CD3^-^ Ly6G^+^), monocytes (CD45^+^ CD3^-^ Ly6G^-^ CD64^hi/low^ Ly6C^+^), macrophages (CD45^+^ CD3^-^ Ly6G^-^ CD64^hi^ Ly6C^-^).

**Table 2.** Flow cytometry and T cell isolation antibodies.

| Target | Clone | Fluorophore | Manufacture | Catalog # |
| --- | --- | --- | --- | --- |
| Ter119 | Ter119 | Biotin | Tonbo Biosciences | 30-5921-U500 |
| CD8a | 53-6.7 | Biotin | Tonbo Biosciences | 30-0081-U500 |
| Cd11b | M1/70 | Biotin | Tonbo Biosciences | 30-0112-U500 |
| Cd11c | N418 | Biotin | Tonbo Biosciences | 30-0114-U500 |
| B220 | RA3-6B2 | Biotin | Tonbo Biosciences | 30-0452-U100 |
| CD317 | eBio927 | Biotin | Invitrogen | 13-3172-82 |
| NK1.1 | PK136 | Biotin | Tonbo Biosciences | 30-5941-U100 |
| CD45 | 30-FF1 | efluor450 | eBioscience | 48-0451-82 |
| CD3 | 145-2C11 | APC | Tonbo Biosciences | 20-0031-U100 |
| CD4 | V4 | AF700 | Biolegend | 100430 |
| CD64 | REA286 | PE | Miltenyi Biotec | 130-118-684 |
| Ly6G | 1A8 | FITC | Tonbo Biosciences | 35-1276-U100 |
| Ly6C | HK1.4 | PE-Cy7 | Invitrogen | 25-5932-82 |
| gp38 | 8.1.1 | AF488 | Biolegend | 127405 |
| CD31 | MEC13.3 | APC/Cyanine7 | Biolegend | 102533 |

### Determining *Tac1* Expression on Sorted Cells

Luria broth (LB)-treated and *C.rodentium* infected WT mice were euthanized at 10 dpi and colons excised. Single-cell suspensions from the lamina propria were isolated as described above. Cells were stained with anti-CD45, anti-CD31, anti-gp38, anti-CD3, anti-Ly6G and Live/Dead Fixable Aqua. Lymphatic endothelial cells (CD45-CD31+ gp38+), blood endothelial cells (CD45-, CD31+, gp38-), T cells (CD45+ CD3+), and neutrophils (CD45+ Ly6G+) were sorted and subsequently processed using the Takara CellAmp Direct TB Green RT-qPCR kit (Cat # 3735A). Intestinal epithelial cells were isolated by longitudinal colectomy and subsequent gentle dissociation in chelating buffer. qPCR was performed using primers specific for *Tac1* obtained from Primerbank. CT values were first normalized to *B-actin* and consequently to dorsal root ganglion, which is known to robustly express *Tac1*.

### Ussing Chambers Assay of Mouse Intestinal Barrier Function

Segments of proximal colon were collected and placed oxygenated Ringer’s buffer (pH 7.35 ± 0.02, with the following composition (in mM): 115 NaCl, 25 NaHCO_3_, 1.2 MgCl_2_, 1.25 CaCl_2_, and 2.0 KH_2_PO_4_). Colonic tissues were opened along the mesenteric border, gently rinsed to remove luminal contents, and mounted in Ussing chambers (Physiologic Instruments, San Diego, CA). Tissues were maintained by continuous oxygenation, at 37 °C with glucose (10 mM) added to the serosal chamber, and an equimolar concentration of mannitol (10 mM) solution added to the mucosal side. Transepithelial short-circuit current (Isc; μA/cm²), potential difference, and conductance (G; mS/cm^2^) were measured after a 15-minute equilibration period until the end of the experiment. Paracellular macromolecular permeability was assessed by the mucosal to serosal flux of 4 kDa. FITC–dextran (Sigma-Aldrich) by sampling the serosal buffer every 30 minutes for 2 hours. Quantification of FITC concentrations in samples was performed using a fluorescence plate reader (excitation: 485 nm; emission: 538 nm) using a standard curve. At the conclusion of permeability measurements, tissues were stimulated with carbachol (CCH, 100 μM), followed by forskolin (20 μM).

### Statistics

Statistical analysis of all data was performed using Prism 10.0 (GraphPad, La Jolla, CA) with a student’s t test or one-way or two-way ANOVA followed by post-hoc analysis with Tukey’s multiple comparison test. Individual data points are presented as mean ± standard error of the mean.

## Author Contributions

EL, MC, MG, and CR designed research studies, EL, MC, KS, and JP conducted experiments, including data acquisition and analysis, EL and CR wrote and edited the manuscript, and CR obtained funding for these studies.

## Funding

These studies were funded by NIH NIAID R01AI150647 (C.R.), Animal Models of Infectious Disease Training Program T32AI060555 (M.C.), and Initiative for Maximizing Student Development (IMSD) Program T32GM135741 (E.L.).

## Results

### Reduced *C. rodentium* shedding and burden in *Tac1^-/-^* mice

Infection with *C. rodentium* resulted in significantly reduced fecal bacterial shedding **(Fig 1A)** and the number of colonic adherent bacteria **(Fig 1B)** in *Tac1^-/-^* mice compared to WT controls 10 days post-infection (dpi). Given the established role of SP in regulating colonic motility, we evaluated distal colonic motor function by measuring fecal output in uninfected mice. As expected, there was a significant reduction in the number **(Fig 1C)** and weight **(Fig 1D)** of fecal pellets in *Tac1^-/-^* compared to WT mice. Although SP can also function as a secretagogue, inducing intestinal ion transport, no significant changes in short circuit current **(Fig S1A),** conductance **(Fig S1B)**, or macromolecular transport indicated by the luminal to serosal translocation of 4 KDa FITC-Dextran **(Fig S1C)** were observed in WT compared to *Tac1^-/-^* mice. We also observed that the physiological ISC response to carbachol was not different in WT versus T*ac1^-/-^ mice* **(Fig S1D)**. Infection-induced colonic histopathology was assessed using H&E-stained colonic tissue sections **(Fig 1E)**, and crypt hyperplasia was assessed. A significant reduction in crypt length was observed in infected *Tac1^-/-^* compared to WT mice **(Fig 1F)**.

**Figure 1.**
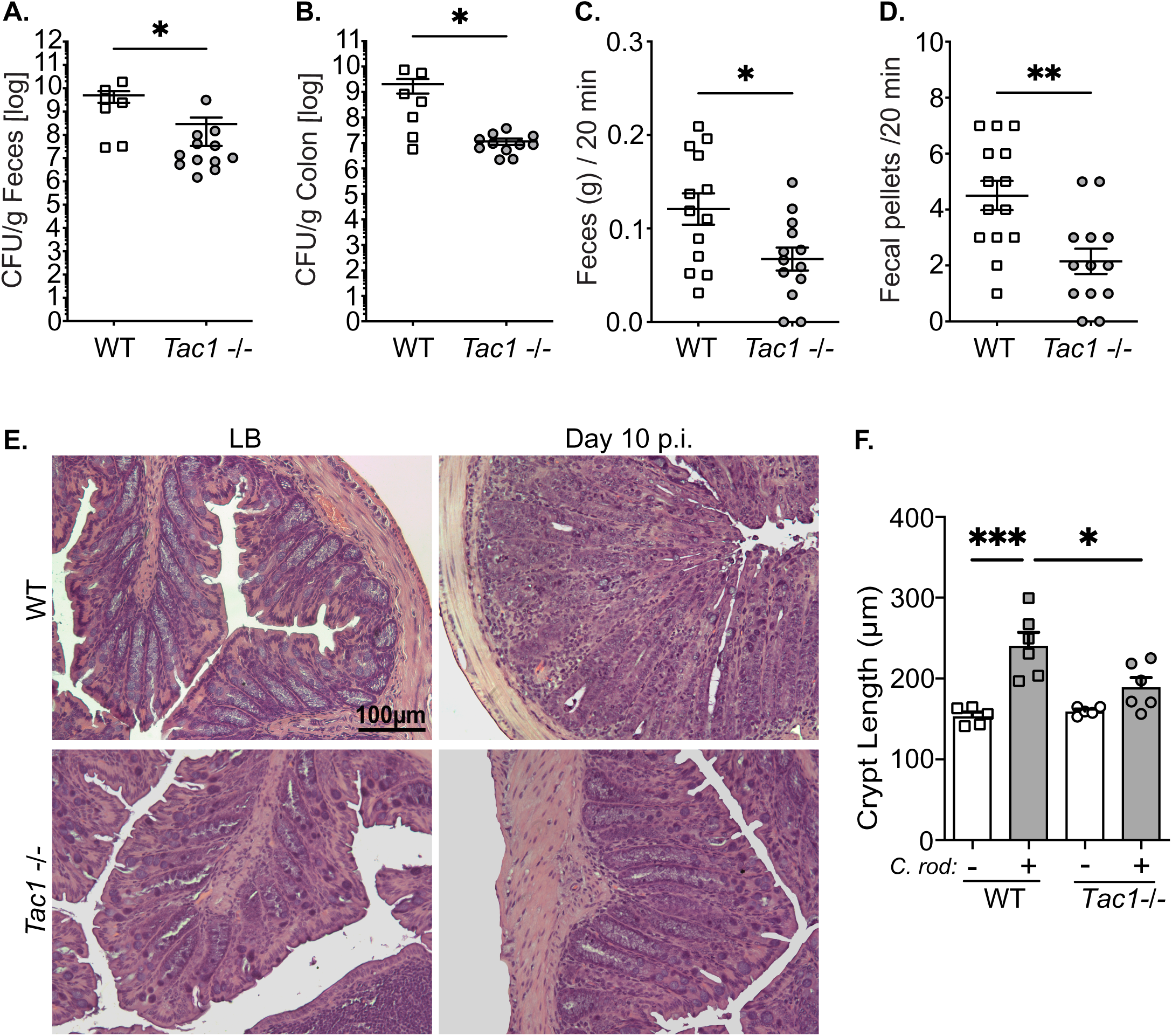
Reduced *C. rodentium* burden and histopathology in Tac1^-/-^ mice. *C. rodentium* bacterial infection in the feces **(A)** and the colonic tissue **(B)** of wildtype (WT) and Tac1^-/-^ mice 10 days post-infection. Distal colonic motility was assessed by the weight of fecal output **(C)** and the number of pellets **(D)**. Colonic histopathology was assessed in H&E sections **(E)**, with quantification of crypt length **(F)**. Data presented from individual mice with mean SD, *P<0.05, ** P< 0.005, ***P<0.001. Student’s two-tailed t-test (A-D), ANOVA followed by post-hoc analysis with Tukey’s multiple comparison test (F). scale bar = 100 µm.

### Tac1-deficient mice exhibit reduced innate immune cell responses to *C. rodentium*

Infection-induced colonic inflammation was assessed in uninfected and infected WT and *Tac1^-/-^* mice. Although *C. rodentium* infection significantly increased expression of the proinflammatory cytokine *Tnfa* in WT mice, this was significantly reduced in *Tac1^-/-^* mice 10 days post-infection **(Fig 2A)**. Despite this, there was no effect of *Tac1* deficiency on the expression of *Il1b* or *Il6 in uninfected or infected mice* **(Fig 2 B&C)**. As expected, *C. rodentium* infection significantly increased expression of *RegIIIγ* irrespective of genotype **(Fig 2D)**. In stark contrast, *Nos2* and *Nox1* expression were significantly upregulated at 10 dpi only in WT, but not *Tac1^-^*^/-^ mice **(Fig 2E & F)**. We further used qPCR to assess cell populations in naïve and *C. rodentium-*infected WT mice with an intact *Tac1* gene to better understand the cellular source of the neuropeptide. Using this approach, we found that compared to the dorsal root ganglia, *Preprotachykinin-1 (Tac1)* expression was exceedingly low in intestinal epithelial cells (IEC), colonic T-cells, neutrophils, as well as blood and lymphatic endothelium **(Fig S2)**.

**Figure 2.**
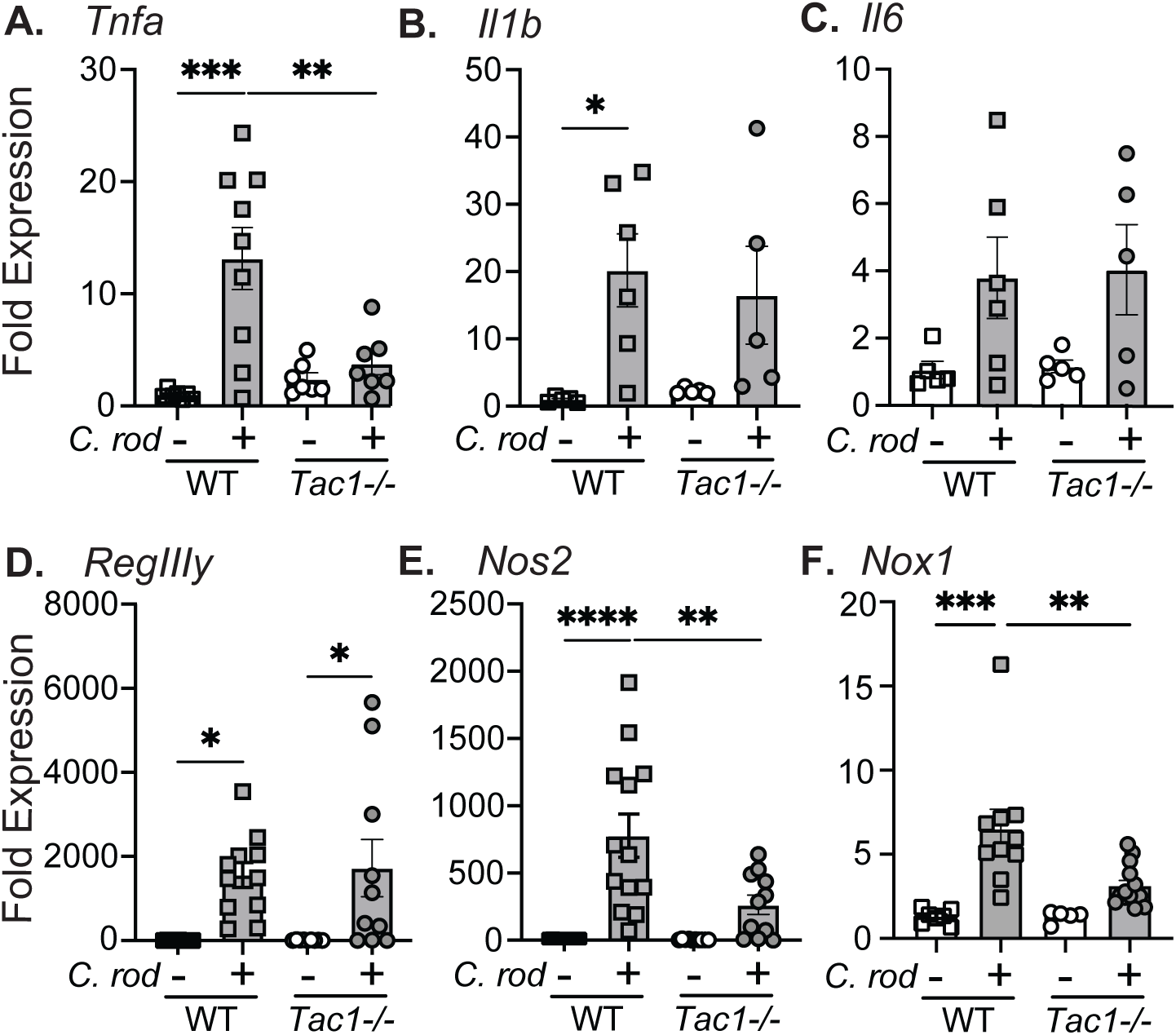
Deficiency in *Tac1-/-* abrogates expression of select host-protective genes. Full-thickness colonic tissues from uninfected control or *C. rodentium* infected WT and *Tac1-/-* mice were assessed for mRNA expression of the proinflammatory cytokines *Tnfa* **(A)**, *Il1b* **(B)**, and *Il6* **(C)**. Expression of *RegIIIy* **(D)**, *Nos2* **(E)**, and *Nox1* **(F)** mRNA was assessed using qRT-PCR. Data are presented from individual mice with mean ± SD, *P<0.05, ** P< 0.005, ***P<0.001, one-way ANOVA followed by post-hoc analysis with Tukey’s multiple comparison test.

Reduced recruitment of innate effector immune cells in *Tac1^-/-^* mice during *C. rodentium* infection.

We sought to further characterize the innate immune response during *C. rodentium* infection in *Tac1^-/-^* compared to WT mice. As neutrophils and monocytes are critical to host protection, we first assessed the expression of chemokines known to regulate homing of these cell populations. Although *Cxcl1* was not reduced significantly, significant reductions in *Cxcl2*, *Cxcl3*, and *Cxcl6* were observed in infected *Tac1^-/-^* compared to WT mice 10 dpi **(Fig 3A-D)**. Although the expression of *Ccl2* increased in WT and *Tac1^-/-^* mice infected with *C. rodentium*, this was not statistically significant across genotypes **(Fig 3E)** suggesting that select chemokines are modulated by infection and deficiency in *Tac1*.

**Figure 3.**
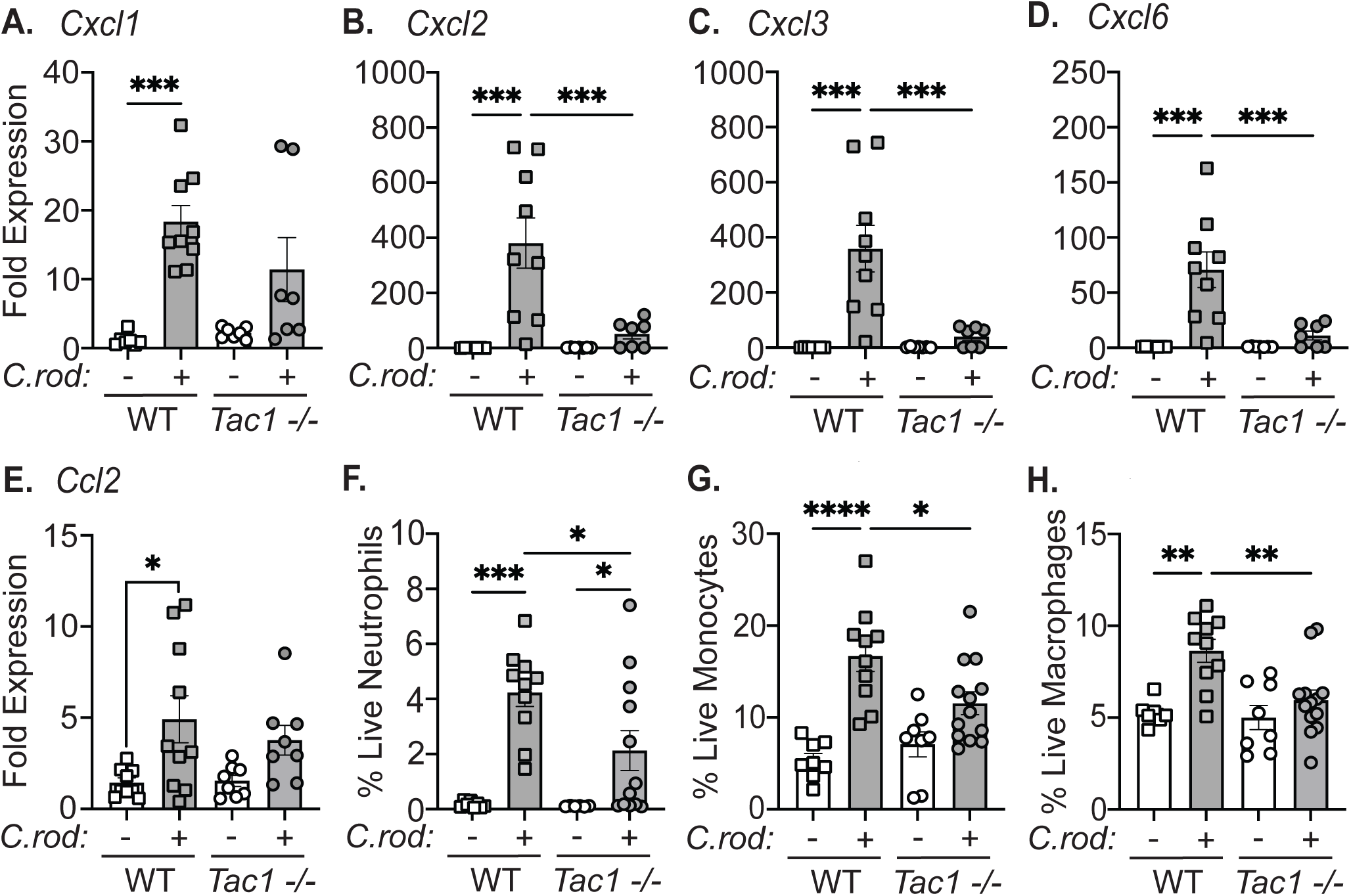
Decreased innate immune cell recruitment in *Tac1-/-* mice during C*. rodentium* infection. Expression of the chemokines, *Cxcl1* **(A)**, *Cxcl2* **(B)**, *Cxcl3* **(C)**, *Cxcl6* **(D)**, *Ccl2* **(E)**, in the colon of uninfected control or *C. rodentium* infected (10 d.p.i.) WT and *Tac1-/-* mice were assessed. Flow cytometry was conducted on single cell suspensions from the colonic lamina propria to determine the frequency of live neutrophils (CD45^+^ CD3^-^ Ly6G^+^) (F), monocytes (CD45^+^ CD3^-^ Ly6G^-^ CD64^hi/low^ Ly6C^+^) (G), and macrophages (CD45^+^ CD3^-^ Ly6G^-^ CD64^hi^ Ly6C^-^) (H). Data presented as individual mice with mean ± SD, *P<0.05, ** P< 0.005, ***P<0.001, one-way ANOVA followed by post-hoc analysis with Tukey’s multiple comparison test.

To understand the effect of these changes in chemokine expression on the recruitment of innate immune cells, we performed immunophenotyping of the colonic lamina propria from uninfected or *C. rodentium*-infected mice 10 dpi (gating strategy, **Fig S3)**. As expected, infection significantly increased the frequency of neutrophils in WT and *Tac1^-/-^* mice compared to uninfected controls. Despite this *Tac1^-/-^* mice exhibited significant reductions in neutrophil recruitment compared to WT controls at the same time point **(Fig 3F)**. Additionally, the frequency of colonic monocytes and macrophages was significantly reduced at 10 dpi in *Tac1^-/-^* compared to WT mice **(Fig 3G)**. These data suggest that *Tac1* gene products regulate innate immune cell responses during *C. rodentium* infection.

### Adaptive immune responses to enteric bacterial infections are reduced in *Tac1-/-* mice

As the host response to *C. rodentium* infection is well established to require coordinated recruitment and function of immune cells, we assessed the effect of *Tac1 deficiency* on adaptive immune cells. *Ifnγ* expression was significantly increased in the colon 10 dpi of WT mice compared to uninfected controls, but this response was significantly diminished in infected *Tac1^-/-^* mice **(Fig 4A)**. Not all cytokine responses were significantly reduced in *Tac1^-/-^* mice compared to WT, as indicated by the expression of *Il17A* and *Il22* observed in infected mice **(Fig 4B&C)**. These changes in IFNγ further prompted us to evaluate *Il12* expression, which was not significantly increased during infection in WT or *Tac1^-/-^* mice **(Fig 4D)**. We further assessed if a reduced ability to produce IFNγ was due to an intrinsic defect in T-cells from *Tac1^-/-^* mice. *In vitro* differentiation revealed no significant difference in the ability to polarize T-cells from the MLN under Th1 conditions **(Fig S3)**. These data suggested that reduced Th1 function of T-cells in the colonic lamina propria of infected *Tac1-/-* mice are not due to a T-cell intrinsic defect.

**Figure 4.**
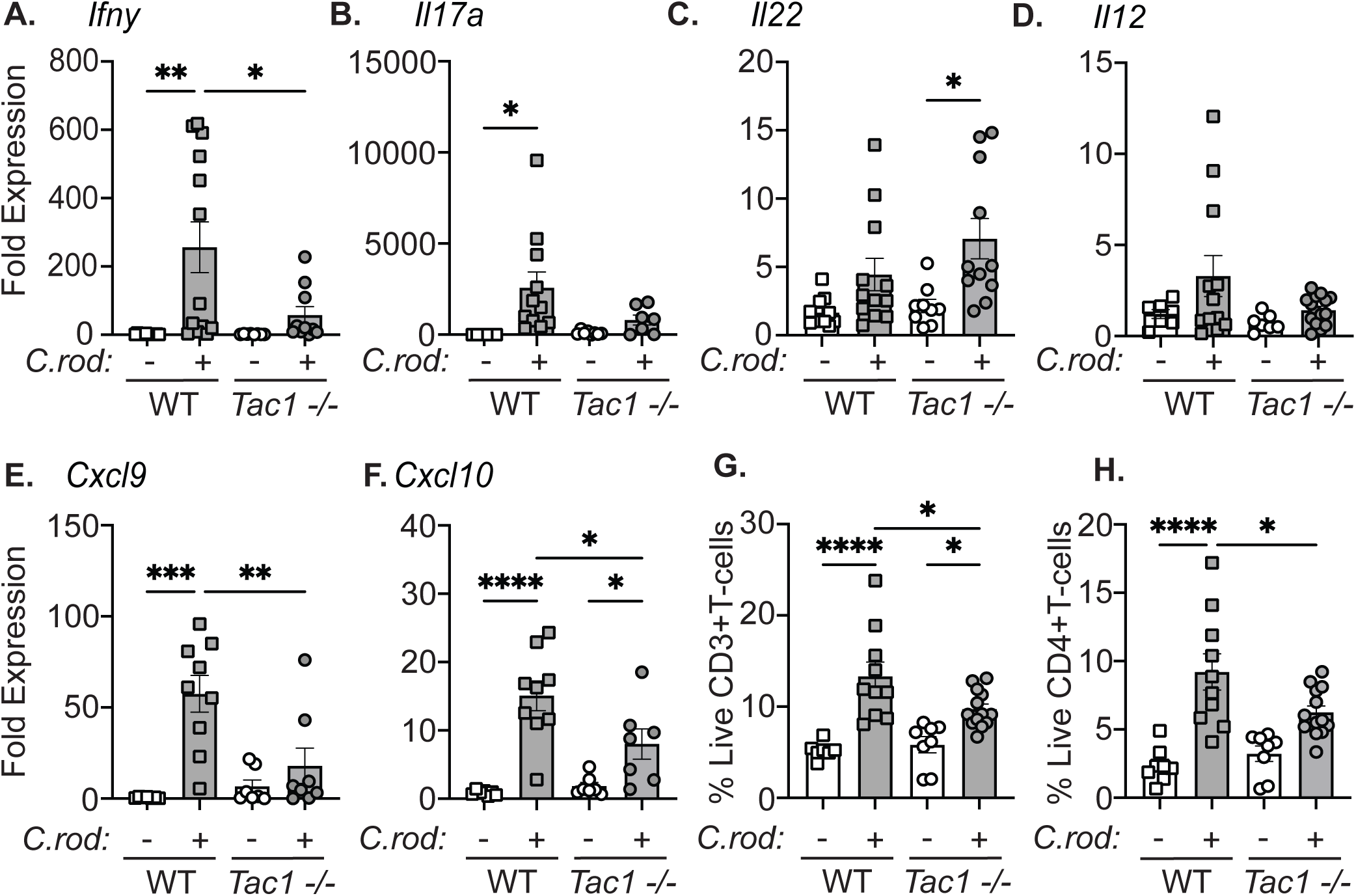
T-cell recruitment is diminished in *C. rodentium*-infected *Tac1^-/-^* mice. Expression of the cytokines *Ifng* **(A)**, *Il17a* **(B)**, *Il22* **(C),** and *Il12* **(D)** mRNA in the colon of uninfected or infected WT and *Tac1-/-* mice was evaluated by qRT-PCR. Quantification of the chemokines *Cxcl9* **(E)**, and *Cxcl10* **(F)** mRNA in the colon of sham or infected WT and *Tac1-/-* mice 10 dpi. Flow cytometry was performed on single-cell suspensions from the colonic lamina propria to identify the frequency of T-cells (live, CD45^+^ CD3^+^), and CD4+ T-cells (live CD45^+^ CD3^+^ CD4^+^). Data are presented from individual mice with mean ± SD, *P<0.05, ** P< 0.005, ***P<0.001, one way ANOVA followed by post-hoc analysis with Tukey’s multiple comparison test.

As chemokines are a critical mechanism to recruit T-cells into the colon during enteric bacterial infection, we assessed the expression of common T-cell chemokines. Significant attenuation of the T-cell chemokines *Cxcl9* and *Cxcl10* were observed in *Tac1^-/-^* versus WT-infected mice **(Fig 4E&F)**. To assess the potential functional implications of reduced chemokine and cytokine expression, flow cytometry was performed to assess T cell recruitment. During *C. rodentium* infection, *Tac1^-/-^* mice had significantly reduced CD4+ T-cells in the lamina propria compared to WT controls **(Fig 4G&H)**. These data suggest that the *Tac1*-encoded gene products are critical for recruiting IFNγ-producing T-cells during enteric bacterial infection.

## Discussion

Mucosal immune responses in the intestinal tract are tightly regulated processes. Although rapid and robust responses are required at this interface to the external environment due to potential pathogen entry, responses to dietary and commensal microbiota-derived substances can be deleterious. Host responses to enteric bacterial pathogens such as *C. rodentium* require successive activation of resident and recruited immune cell populations. Effective coordination of these processes occurs through numerous overlapping mechanisms, including bidirectional communication between the nervous and immune systems. These systems are interconnected through the expression of neurotransmitter receptors by numerous types of immune cells, coupled with the ability of neurons to detect pathogens and immune cell-derived products. These include the detection of cytokines by specialized sensory neurons within the affected tissues of immune organs [9, 21]. During *C. rodentium* infection, the polymodal nociceptive protein TRPV1 that is expressed by these sensory neurons plays a critical role in the recruitment of neutrophils to the infected colon [8]. It is well appreciated that these neurons can release neuropeptides such as Substance P locally when activated by noxious stimuli, resulting in neurogenic inflammation [22]. The molecular underpinnings of this response are due to activation of Substance P receptor on a diverse array of cell types, including endothelial cells, to induce vascular permeability and increase cellular adhesion molecule expression. We have previously demonstrated that the Substance P receptor has a critical role in the regulation of colonic inflammation during *C. rodentium* infection [10]. Complicating the interpretation of this role of Substance P receptor signaling in mediating host protection, other ligands such as hemokinin-1 (aka Tac4) can activate the SP receptor [23]. With this complexity in mind, we sought to provide a more comprehensive picture of the role of the neuropeptides encoded by *Tac1* in regulating host defense. Here, we have demonstrated that select aspects of the immune response to *C. rodentium* are altered in *Tac1-*deficient mice. Infection of *Tac1^-/-^* mice with *C. rodentium* significantly reduced bacterial burden and fecal shedding compared to WT mice. These data were in keeping with our prior studies using pharmacological antagonists of the substance P receptor [10]. Given the known roles of Substance P as a positive regulator of colonic motility [24], we considered whether fecal pellet output as a measure of distal colonic motility was altered in *Tac1^-/-^* compared to WT mice. Demonstrating that the observed reduced bacterial burden in *Tac1^-/-^* mice was not due to increased colonic motility, we observed significantly reduced fecal pellet output compared to WT controls. These data are also in keeping with observations of reduced fecal output with Substance P receptor antagonism [10]. As Substance P has long been described as secretagogues that increase the secretory state of the gastrointestinal tract [25, 26], we performed Ussing chamber studies with colonic tissues from WT and *Tac1^-/-^* mice. We noted no significant differences in the baseline or evoked Isc, conductance, or macromolecular permeability in *Tac1^-/-^,* suggesting differences in host immune responses were not a consequence of an altered baseline intestinal physiology.

As the immune response to *C. rodentium* is well-appreciated to induce colonic epithelial cell hyperplasia, we assessed this host response to infection. Reduced bacterial burden in *Tac1^-/-^* mice compared to WT was associated with reduced colonic crypt depth and hyperplasia. In WT mice, these structural changes are a consequence of inflammation and pro-inflammatory cytokines increasing epithelial cell proliferation and the number of non-differentiated epithelial cells along the crypt axis [27]. Infection-induced increases in IFNγ-IFNγ receptor signaling are a known driver of crypt hyperplasia, as infection of IFNγ receptor-deficient mice failed to induce crypt hyperplasia [28, 29]. Our data showing reduced expression of pro-inflammatory cytokines in infected *Tac1^-/-^* mice suggests that reduced histopathology is due to reduced inflammation. Here we demonstrate that *Tac1^-/-^* mice have significantly reduced IFNγ expression and corresponding reductions in IFNγ-regulated genes, including *Nos2* [30] and *Nox1*, in *C. rodentium*-infected *Tac1^-/-^* vs. WT mice. While the enzymes encoded by these genes are typically regarded as host protective, *Nos2^-/-^* mice have reduced bacterial clearance without increased morbidity or mortality during *C. rodentium* infection [30, 31]. More recently, the production of H_2_O_2_ has also been shown to be host maladaptive, supporting the proliferation of enteric bacterial pathogens, including *C. rodentium* [32]. Although elevated colonic nitrate due to increased expression and activity of Nos2 supports the growth of *Salmonella enterica* and select commensal bacteria, *C. rodentium* does not use nitrate as a terminal electron acceptor in support of metabolism and growth. However, inflammation and the ensuing colonic oxygenation increases *C. rodentium* proliferation [33]. These data suggest that the immune response to *C. rodentium* and colonic inflammation can be host maladaptive. Reduced IFNγ and inflammation in *Tac1^-/-^* mice therefore suggest that, in WT mice, these neuropeptides enhance inflammatory processes that can support pathogen growth. These data are in keeping with effects observed with our prior observations, where antagonism of the substance P receptor reduced bacterial burden, IFNγ, and IFNγ-dependent gene expression during *C. rodentium* infection. Together, these data suggest the enhancement of IFNγ production by a SP-SP receptor signaling axis during enteric bacterial infection, and that this response can evoke host maladaptive responses. Also of note, we observed reduced *Tnfa* in infected *Tac1^-/-^* mice, a feature not observed in our prior SP receptor antagonists studies [10]. These results may suggest a possible role for neurokinin A, as the other peptide encoded by the mouse *Tac1* gene, or that an alternative receptor could be engaged. Substance P can bind and activate the Mas-related G protein-coupled receptors (Mrgprs) to induce a variety of biological effects ranging from activation of nociceptors to causing degranulation of mast cells [34, 35]. While mast cells have been demonstrated to exert an antibacterial effect in *C. rodentium* infection [36], and Mrgprs-mediated activation of mast cells limits skin infection [30], it is unknown if SP-induced Mrgprs signaling can affect mast cell-dependent host responses.

Substance P exerts additional control over the immune response during infection, with Tac1-deficient mice exhibiting a clear reduction in the expression of select chemokines that result in neutrophil and T-cell recruitment. Significant reductions in these chemokines were further matched by reduced recruitment of these cell types to the infected colon of *Tac1^-/-^* mice compared to WT. Once again, these reductions in T-cell recruitment mirror the effects observed in infected Substance P receptor antagonist-treated mice that we reported previously [10]. These findings suggest that Substance P receptor signaling enhances the recruitment of immune cell populations through different mechanisms, each of which could be targeted for modulation. For example, Substance P-induced activation of blood endothelial cells would have different effects on the development of an immune response compared to activation of stromal cells. As part of the unique biological outcomes, not only are the responding cell populations important, but the source of Substance P, and thus the location of the rapidly clearing peptide, is a critical factor to consider as well. Although many sources of Substance P in the intestine have been previously identified, including immune cells, stroma, and intestinal epithelial cells [17, 22, 37], we found low levels *Tac1* expression compared to sensory neurons of the dorsal root ganglia in these cell types in uninfected and *C. rodentium-infected* mice. It is important to note that these data do not exclude the potential for rare but biologically significant sources of Substance P in these populations, only in the bulk population or each cell type there is very low abundance of expression. Indeed, this would likely include the reported Substance P-producing enteroendocrine cells within the colonic IEC [17]. The cell types that produce biologically relevant amounts of SP during *C. rodentium* infection remains an open question that should be addressed using conditional KO mice in future studies. Together, our data are in keeping with the literature on Substance P receptor signaling, demonstrating a significant role of this neuropeptide in the host response during enteric bacterial infection. We further speculate that modulation of this signaling axis may provide a previously unappreciated mechanism to limit maladaptive inflammation without impinging upon bacterial clearance.

## Notes

### Competing Interest Statement

The authors have declared no competing interest.

