## Supplementary figures and images for "Tac1 Deficiency Reduces the Severity of Enteric Bacterial Infection"

### Supplemental figures

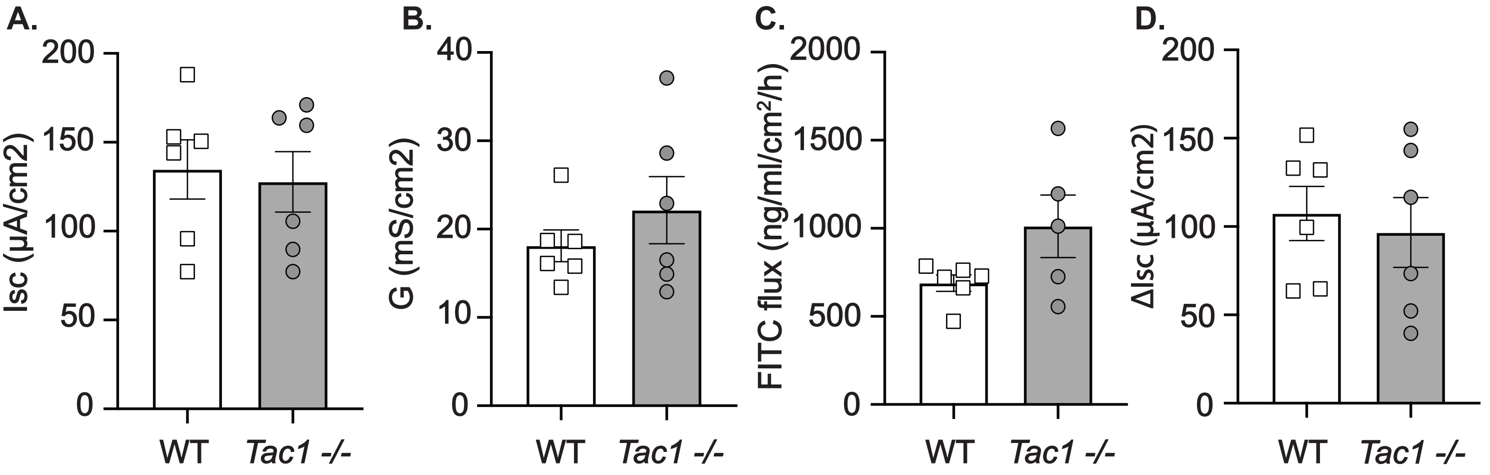

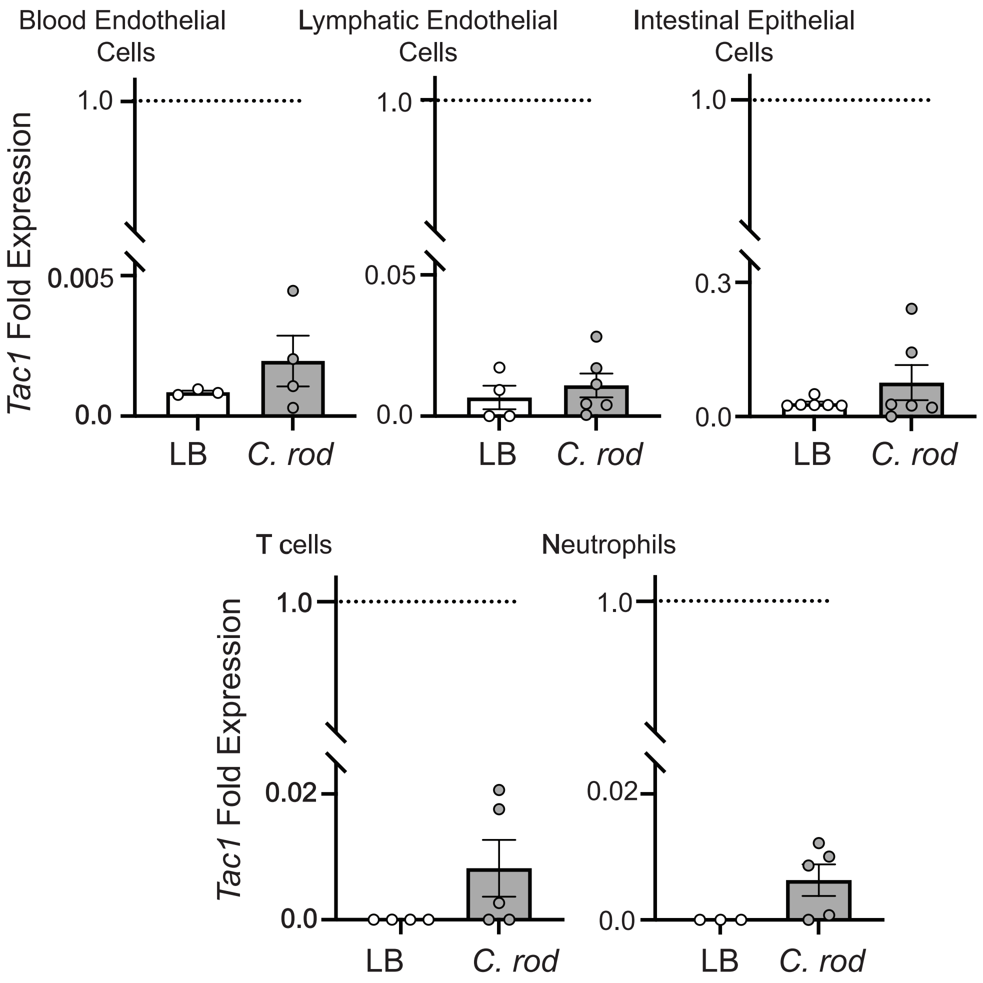

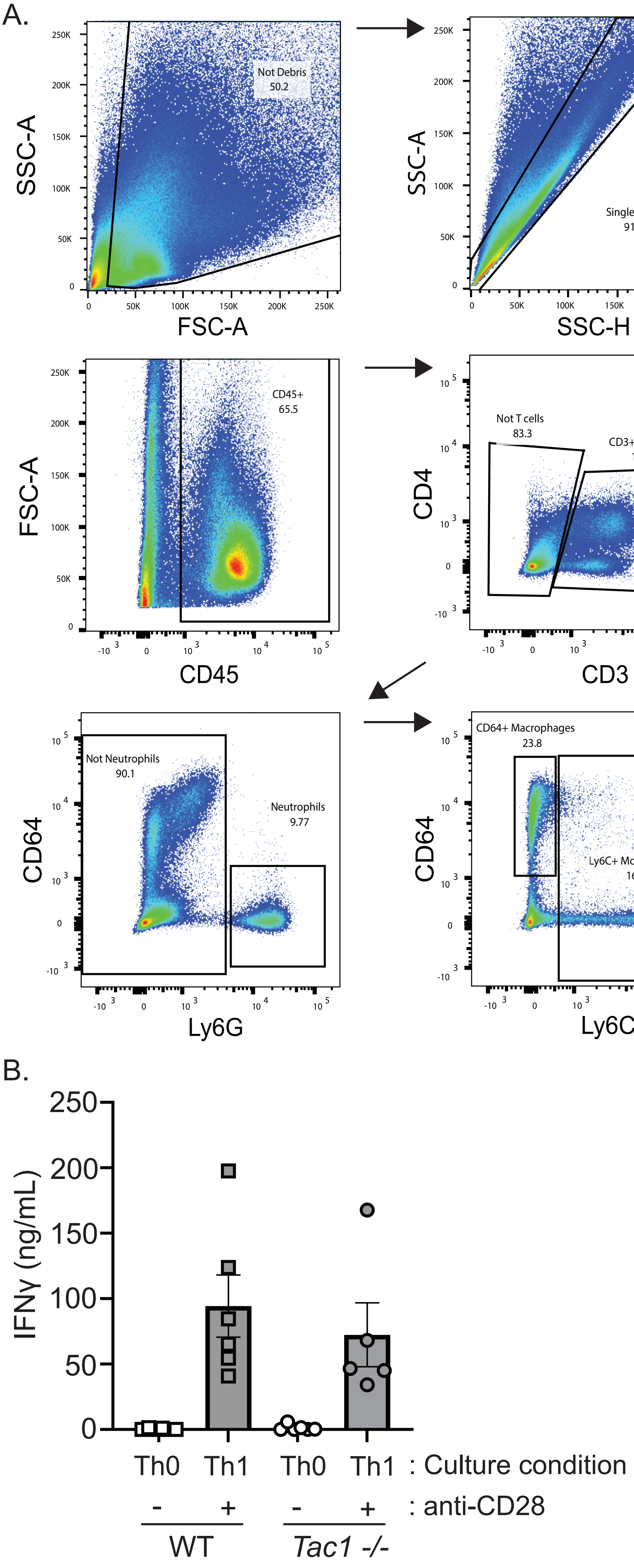
